# Neurodegenerative disease-associated protein aggregates are poor inducers of the heat shock response in neuronal-like cells

**DOI:** 10.1101/2020.01.06.896654

**Authors:** R. San Gil, D. Cox, L. McAlary, T. Berg, A. K. Walker, J. J. Yerbury, L. Ooi, H. Ecroyd

## Abstract

Protein aggregation that results in the formation of inclusions is strongly correlated with neuronal death and is a pathological hallmark common to many neurodegenerative diseases, including amyotrophic lateral sclerosis (ALS) and Huntington’s disease. Cells are thought to dramatically up-regulate the levels of heat shock proteins during periods of cellular stress via induction of the heat shock response (HSR). Heat shock proteins are well-characterised molecular chaperones that interact with aggregation-prone proteins to either stabilise, refold, or traffic protein for degradation. The reason why heat shock proteins are unable to maintain the solubility of particular proteins in neurodegenerative disease is unknown. We sought to determine whether neurodegenerative disease-associated protein aggregates can induce the HSR. Here, we generated a neuroblastoma cell line that expresses a fluorescent reporter under conditions of HSR induction, for example heat shock. Using these cells, we show that the HSR is not induced by exogenous treatment with aggregated forms of Parkinson’s disease-associated *α*-synuclein or the ALS-associated G93A mutant of superoxide dismutase-1 (SOD1^G93A^). Furthermore, flow cytometric analysis revealed that intracellular expression of SOD1^G93A^ or a pathogenic form of polyQ-expanded huntingtin (Htt^72Q^), similarly, results in no or low induction of the HSR. In contrast, expression of a non-pathogenic but aggregation-prone form of firefly luciferase (Fluc) did induce an HSR in a significantly greater proportion of cells. Finally, we show that HSR induction is dependent on the intracellular levels of the aggregation-prone proteins, but the pathogenic proteins (SOD1^G93A^ and Htt^72Q^) elicit a significantly lower HSR compared to the non-pathogenic proteins (Fluc). These results suggest that pathogenic proteins either evade detection or impair induction of the HSR in neuronal-like cells. Therefore, defective HSR induction may facilitate the initiation of protein aggregation leading to inclusion formation in neurodegenerative diseases.

## Introduction

The formation of intracellular protein inclusions is a characteristic hallmark of neurodegenerative diseases, such as amyotrophic lateral sclerosis (ALS), Parkinson’s disease (PD) and Huntington’s disease (HD) (Chiti and Dobson 2017). In these diseases, proteins are either intrinsically disordered or partially unfold and self-associate through hydrophobic interactions between regions that are usually buried in the native conformation (Hipp, Kasturi et al. 2019). These inappropriate interactions nucleate the formation of protein aggregates that are the building blocks of protein inclusions (Kopito 2000). Inclusion formation has been strongly correlated with the death of neurons in neurodegenerative diseases (Braak, Del Tredici et al. 2003, Braak, Alafuzoff et al. 2006, Brettschneider, Del Tredici et al. 2013, Brettschneider, Arai et al. 2014). The progression of protein inclusion pathology from a focal site of onset to other regions of the central nervous system (CNS) may be driven by the movement of aggregated proteins in the extracellular space (Vaquer-Alicea and Diamond 2019). Therefore, aggregation is seeded in neighbouring cells and neurons via prion-like propagation [for reviews see (Jucker and Walker 2013, Zeineddine and Yerbury 2015, Hanspal, Dobson et al. 2017, Victoria and Zurzolo 2017)]. These stages of disease onset and progression throughout the CNS suggest a widespread inability of cells to prevent the initial events leading to protein aggregation and the propagative seeding events that follow.

Cells have several mechanisms of defense against disturbances in protein homeostasis (proteostasis) (Yerbury, Ooi et al. 2016). These mechanisms usually function to ensure the correct folding, function, and turnover of proteins in the cell, and prevent the negative effects of proteostasis imbalance on cell viability. The heat shock response (HSR) is one such mechanism of proteostasis and acts as a first line of defense against protein destabilisation, misfolding, and aggregation (San Gil, Ooi et al. 2017). Increasing intracellular abundance of misfolded proteins can activate heat shock transcription factor 1 (HSF1), which translocates into the nucleus and binds heat shock elements (HSEs; pentameric sequence nGAAn where ‘n’ is any nucleotide) in the promoter elements of its target genes (Pelham 1982, Sorger, Lewis et al. 1987, Amin, Ananthan et al. 1988, Xiao and Lis 1988). Induction of the HSR results in rapid and dramatic upregulation of a family of proteins called heat shock proteins (Hsps), which are well-characterised molecular chaperones capable of stabilising and re-folding misfolded proteins, with additional functions in trafficking damaged proteins to proteasomal or autophagy-lysosome degradation pathways (Leak 2014). Heat shock proteins have previously been shown to interact with disease-associated mature aggregates *in vitro* (Cox, Whiten et al. 2018, Wu, Vonk et al. 2019) and co-localise with pathogenic inclusions in brains of patients with neurodegenerative diseases (Watanabe, Dykes-Hoberg et al. 2001). Together, these findings suggest that Hsps act directly on misfolded proteins and increasing Hsp levels and their activity may be beneficial at both the early and late stages of the process of protein aggregation.

There is a wealth of evidence demonstrating that the over-expression of individual Hsps in cell-based and *in vivo* models significantly ameliorates neurodegenerative disease-associated protein aggregation (Bose and Cho 2017, San Gil, Ooi et al. 2017, Webster, Darling et al. 2019). For example, over-expression of Hsp40 and Hsp27 results in a 90-95% decrease in insoluble protein in QBI-293 (human embryonic kidney) cells expressing an aggregation-prone, acetylation-mimicking mutant of ALS-associated TAR DNA-binding protein-43 (TDP-43) (Wang, Wander et al. 2017). Likewise, over-expression of HSF1 in a mutant superoxide dismutase-1 (SOD1) mouse model of ALS, led to a 34% decrease in the level of insoluble SOD1^H46R/H48Q^ in spinal cord tissue compared to controls (Lin, Simon et al. 2013). The over-expression of constitutively active HSF1 in the R6/2 mouse model of HD led to a 79% reduction in the proportion of nuclei with inclusions and extended survival by 16 days (mean survival 122 days in R6/2-HSF1Tg compared to 102 days in R6/2 mice) (Fujimoto, Takaki et al. 2005). Therefore, activating the HSR and over-expressing Hsps in the CNS has promising therapeutic potential to ameliorate protein aggregation and extend survival in neurodegenerative diseases.

The activation of HSF1 may be differentially stimulated by misfolded proteins depending on whether they are sequestered into iPODs (Insoluble PrOtein Deposits), JUNQ (JUxtaNuclear Quality control), or other undefined sub-types of protein inclusions (Kaganovich, Kopito et al. 2008). There is compelling evidence to suggest that HD-associated poly-glutamine expanded huntingtin is sequestered into iPODs, immobile inclusions for the storage of terminally aggregated proteins, that have little interaction with the proteostasis network (Kayatekin, Matlack et al. 2014, Polling, Mok et al. 2014). In contrast, it has been demonstrated that mutant SOD1 is partitioned into JUNQ compartments in which proteins are able to diffuse and are ubiquitinated, indicating a role for the ubiquitin-proteasome system (Matsumoto, Kim et al. 2006, Polling, Mok et al. 2014, Farrawell, Lambert-Smith et al. 2015). In addition to the sub-type of inclusion formed, the propensity of different neurodegenerative disease-associated proteins to undertake different off-folding pathways, *i.e.* amorphous or amyloid aggregation, may represent another factor that determines whether an HSR is induced.

The anti-aggregation and cytoprotective benefits of Hsps in cells cannot be realised if an HSR is not induced in neurons and glia in response to protein aggregation in neurodegenerative disease. The existence of intracellular inclusions and the cell-to-cell spread of aggregated proteins in neurodegenerative diseases led us to hypothesise that cells are not able to respond to protein aggregation by inducing the HSR. To evaluate this, we generated a stable cell line derived from mouse neuroblastoma, Neuro-2a, that express a fluorescent reporter under the control of a truncated *Hspa1a* promoter comprised of 8 putative HSEs. We then used this stable cell line to quantitatively assess the kinetics and magnitude of HSR induction after exogenous application of protein aggregates and intracellular inclusion formation. Moreover, we dissected the role of inclusion size, rate of formation, and intracellular protein concentration on HSR induction using these cells. We identified that whilst cells were able to activate the HSR in response to aggregation-prone non-pathogenic proteins, they failed to mount a significant HSR in response to the exogenous application or intracellular expression of pathogenic proteins. In addition, the intracellular levels of all aggregation-prone proteins tested was the most significant determinant of HSR induction and demonstrated a positive correlation between protein concentration and proportion of cells with an activated HSR. The relatively poor induction of the HSR by pathogenic compared to non-pathogenic proteins could explain, at least in part, the ability of aggregates to evade molecular chaperones, form protein inclusions, and subsequently spread to other regions of the CNS in the context of neurodegenerative diseases.

## Materials and methods

All reagents and chemicals used in this work were obtained from Merck-Sigma Aldrich (NSW, Australia) and Amresco (OH, USA) unless otherwise stated.

### Plasmids

To generate cell lines that stably and constitutively express mCherry and stress-inducible EGFP downstream of a minimal Hsp70 promoter (minHsp70p) the following two constructs were generated; pCMV-mCherry and pminHsp70p-EGFP. With respect to pCMV-mCherry, the EGFP gene was excised from pEGFP-N1 (Takara Clontech, France) with flanking *Eco*RI/*Bsr*GI sites and replaced with the mCherry gene. Regarding pminHsp70p-EGFP, a minimal Hsp70 promoter consisting of 8 putative heat shock elements (HSE; conserved pentameric sequence; nGAAn) upstream of an EGFP gene was excised with flanking *Acc*651/*Bam*HI sites (kind gift of Dr Franck Couillaud, University of Bordeaux, France, and Dr Chrit Moonen, UMC Utrecht, The Netherlands) and subcloned into *Acc*651/*Bam*HI digested pGL4.4 (Thermo Fisher Scientific, VIC, Australia) containing a hygromycin resistance gene.

The constructs coding for the expression of Cerulean-tagged huntingtin exon 1 fragment (Htt) with a non-pathogenic (i.e. 25 polyglutamines; pT-Rex-Cerulean-Htt^25Q^) or pathogenic (i.e. 72 polyglutamines; pT-Rex-Cerulean-Htt^72Q^) poly-glutamine(Q) tracts were kind gifts from Prof Danny Hatters (University of Melbourne, VIC, Australia). Constructs for the expression of Cerulean-tagged SOD1^WT^, SOD1^G93A^, WT firefly luciferase (Fluc^WT^) and a double mutant (R188Q/R261Q) form of Fluc (Fluc^DM^) were also generated. To do so, the *cerulean* gene was PCR amplified from pT-Rex-Htt^72Q^ using forward: 5′-catggatccaccggtcgccaccatggtgagca-3′ and reverse: 5′-caggattcttacttgtacagctc-3′ primers with flanking *Bam*HI/*Bsr*GI restriction sites to replace the EGFP gene in pEGFP-N1-SOD1^WT^ and pEGFP-N1-SOD1^G93A^. The *cerulean* gene was PCR amplified from pT-Rex-Htt^72Q^ using forward 5′-catgggatccaccggccggtcgccaccatggtgagc-3′ and reverse: 5′-caggattcttacttgtacagctc-3′ primers with flanking *Bam*HI/*Bsr*GI restriction sites to replace the EGFP gene in pcDNA4-TO-myc-hisA-EGFP-Fluc^WT^ and pcDNA4-TO-myc-hisA-EGFP-Fluc^DM^ (these constructs were kind gifts of Prof Mark Wilson, University of Wollongong, NSW, Australia). All the constructs generated and used in this work were verified by sequencing using a Hitachi 3130xl Genetic Analyser (Applied Biosystems, MA, USA).

### Generation of SOD1^G93A^ and α-synuclein aggregates

#### Thioflavin T-based aggregation assays

The formation of SOD1^G93A^ aggregates was monitored by an *in situ* thioflavin T (ThT) binding assay that has previously been described (McAlary, Aquilina et al. 2016). Briefly, 100 µM purified SOD1^G93A^ was incubated with 20 mM DTT, 5 mM EDTA and 10 µM ThT in PBS (pH 7.4) at 37°C. The reaction mixtures were loaded into a clear-bottomed 384-well plate (Greiner, Germany). The plate was incubated in a PolarStar Omega Plate Reader (BMG Labtechnologies, VIC, Australia) at 37°C for 30 min prior to covering with adhesive film and commencing readings of samples. The plate underwent double orbital shaking at 300 rpm for 300 s at the start of a 900 s cycle for at least 200 cycles. The ThT fluorescence was measured by excitation at 450 nm and its emission read at 480 nm using the bottom optic.

The formation of α-synuclein fibrils was determined by an end-point ThT assay as previously described (Buell, Galvagnion et al. 2014). Briefly, α-synuclein seeds were produced by incubating 150 µM α-synuclein in 50 mM phosphate buffer (pH 7.4), for 48 h at 40°C under maximal stirring with a magnetic stirrer (WiseStir MSH-20A, Witeg, Germany). The seed fibrils were fragmented by sonication using a microtip probe sonicator (Branson 250 Digital Sonifer, Branson Ultrasonics, CT, USA), using 30% amplification and 3 cycles of 10 s pulses. The fibrils were flash frozen in liquid nitrogen and stored at -80°C until required. To produce mature fibrils, 100 µM of monomeric α-synuclein was incubated with 1% (w/w) α-synuclein seeds in 50 mM phosphate buffer (pH 7.4) at 37°C for 72 h. A 5 µL aliquot of the α-synuclein fibrils was then loaded into a black clear-bottomed 384-well plate with 25 µL 50 mM phosphate buffer containing 10 µM ThT. The ThT fluorescence was measured by excitation at 450 nm and its emission read at 480 nm using the bottom optic.

#### Transmission electron microscopy of SOD1^G93A^ and α-synuclein

Transmission electron microscopy (TEM) was employed to visualise the recombinant SOD1^G93A^ and α-synuclein aggregates. An 8 µL aliquot of the aggregated protein was applied onto an ultrathin carbon film coated 400 mesh copper TEM grid (ProSciTech, QLD, Australia). Samples were then diluted with 2 µL of 0.22 µm filtered milli-Q H_2_O and left for 3 min. Grids were dried with lint-free paper by wicking away the H_2_O from the side. Grids were washed with 10 µL of milli-Q H_2_O, dried again, and 10 µL of 1% (w/v) phosphotungstic acid, the contrast reagent, was added and grids left to incubate for 1 min. The phosphotungstic acid was wicked away and the grids were washed twice with 10 µL of milli-Q H_2_O. Grids were air dried and imaged at the Australian Institute of Innovative Materials (University of Wollongong) using a JEM-2011 TEM (JEOL, Japan). Images were processed using Digital Micrograph (Gatan, CA, USA).

### Cell culture of Neuro-2a

The murine neuroblastoma cell line, Neuro-2a, was obtained from the American Type Culture Collection (VA, USA). All cell lines were cultured in DMEM/F-12 supplemented with 2.5 mM L-glutamine and 10% (v/v) FCS (10% FCS-DMEM/F-12) at 37°C under 5% CO_2_/95% air in a Heracell 150i CO_2_ incubator (Thermo Fisher Scientific). Cells were passaged every 2 days or once they had reached 80% confluency. The cells were tested for mycoplasma on receipt of the cell line and quarterly thereafter using the MycoAlert Mycoplasma Detection Kit according to the manufacturer’s instructions (Lonza, Basel, Switzerland).

### Generation and maintenance of Neuro-2a stable cells

Stable cell lines for the constitutive expression of mCherry and stress-inducible expression of EGFP were generated in Neuro-2a cells. Thus, activation of the HSR (*i.e.* HSF1 binding to HSEs) after treatment with a stressor could be monitored in cells in real time using EGFP as a fluorescent reporter. Neuro-2a cells were used because they have a neuronal origin and high transfection and co-transfection efficiencies using standard lipid-based protocols.

Neuro-2a were first transfected with LTX Plus (Life Technologies, VIC, Australia; 1 µg DNA, 1 µL PLUS reagent, and 3 µL Lipofectamine LTX per well) with *Vsp*I linearised pCMV-mCherry and grown under selective pressure (300 µg/mL G418) for 7 days. Monoclonal mCherry-expressing Neuro-2a cell lines were generated by limiting dilution and subsequent monoclonal expansion. Monoclonal mCherry-expressing Neuro-2a cell lines were transfected with *Not*I linearised pminHsp70p-EGFP and transfected cells were grown under selective pressure (300 µg/mL G418 and 100 µg/mL hygromycin) for 7 days.

To obtain a polyclonal Neuro-2a cell population with stress-inducible EGFP expression, cells were heat shocked (42°C for 2 h with recovery at 37°C for 6 h), harvested by trypsinisation with 0.25% (w/v) trypsin-EDTA at 37°C for 5 min, washed twice in PBS, and resuspended in FACS buffer, 1 mM EDTA, 25 mM HEPES and 0.5% (w/v) BSA in PBS. These cells were then sorted using an S3e Cell Sorter (Bio-Rad Laboratories, NSW, Australia) equipped with 488 nm and 561 nm lasers. Viable, single cells were resolved based on plots of forward scatter-area versus side scatter-area and forward scatter-area versus forward scatter-height. Subsequently, mCherry^+ve^/EGFP^+ve^ cells were identified and sorted and maintained in complete medium supplemented with 1 *×* penicillin/streptomycin to prevent bacterial contamination. These EGFP HSR reporter cell lines are referred to in this work as Neuro-2a (HSE:EGFP), given EGFP expression is driven by HSF1 binding to HSEs.

Neuro-2a (HSE:EGFP) cells were maintained under the same conditions as the parental cell lines. Constant selection pressure was achieved by supplementing the media used to culture Neuro-2a (HSE:EGFP) with 300 µg/mL G418 and 100 µg/mL hygromycin.

### Cell stress treatments

#### Heat shock, cadmium chloride (CdCl_2_), and celastrol treatment assays

Heat shock, CdCl_2_ and celastrol (AdooQ Biosciences, CA, USA) were used to assess the capacity of Neuro-2a cells to induce an HSR. Neuro-2a (HSE:EGFP) cells were seeded at a density of 200,000 cells/mL into 12-well plates. Optimal concentrations of CdCl_2_ and celastrol were determined in concentration-response assays, whereby cells were either treated with CdCl_2_ (0-33 µM) or celastrol (0 – 1 µM) for 24 h. In addition, cells were heat shocked (42°C for 2 h) and allowed to recover for different times at 37°C. After each treatment, Neuro-2a (HSE:EGFP) cells were imaged every 2 h for 24 h using an IncuCyte Live Cell Analysis System (Essen BioScience, MI, USA). The optimal treatment concentrations and times for induction of an HSR in these cells were determined to be 10 µM for 24 h for CdCl_2_, 0.75 µM for 24 h for celastrol and heat shock at 42°C for 2 h with recovery at 37°C for 24 h.

As a means of assessing magnitude and kinetics of HSR induction following each treatment, the maximum EGFP fluorescence intensity and time taken to reach half of the EGFP maximum intensity was analysed.

### Extracellular aggregation stress assays

Pathogenic protein aggregates were applied extracellularly to Neuro-2a (HSE:EGFP) cells and activation of the HSR was assessed. Prior to treatment, soluble non-aggregated α-synuclein and SOD1^G93A^ were centrifuged (14,000 × *g* for 30 min at 4°C) to remove any oligomeric seeds that may have spontaneously formed. Aggregated SOD1^G93A^ was pelleted (14,000 × *g* for 30 min at 4°C) and resuspended in fresh PBS to eliminate possible cytotoxicity of DTT and EDTA in the assay.

Neuro-2a (HSE:EGFP) cells were seeded at a density of 100,000 cells/mL in a 96-well plate and cultured overnight in 10% FCS DMEM/F12. The following day, media was refreshed with serum free DMEM/F12 and cells were either treated with buffer alone (50 mM phosphate buffer for α-synuclein or PBS for SOD1^G93A^), monomeric α-synuclein or aggregated α-synuclein (1 µM and 10 µM), or dimeric SOD1^G93A^ or aggregated SOD1^G93A^ (1 µM and 10 µM) diluted in serum-free DMEM/F12. Three wells in each plate were treated with 10 µM CdCl_2_ as a positive control for HSR induction. Cells were imaged every 2 h for 72 h in an IncuCyte Live Cell Analysis System (Essen BioScience).

### Intracellular protein aggregation stress assays

Neuro-2a (HSE:EGFP) cells were transfected with Cerulean-tagged WT and aggregation-prone mutant proteins. Cells were seeded at a density of 100,000 cells/mL in 12-well plates and cultured in 1 mL of 10% FCS DMEM/F12 media overnight. Cells were transfected with DNA:lipid complexes (1 µg DNA, 1 µL PLUS reagent, and 3 µL Lipofectamine LTX per well) for the expression of Cerulean-tagged SOD1^WT^, SOD1^G93A^, Htt^25Q^, Htt^72Q^, Fluc^WT^, or Fluc^DM^.

As controls, parental Neuro-2a were either untransfected, or singly transfected to express EGFP, mCherry or Cerulean fluorescent proteins. These samples were used to set gates for the flow cytometric analysis and to determine the spectral overlap that occurs between these three fluorophores so that spectral compensation could be applied prior to analysis. All analyses of the flow cytometry data were performed using FlowJo (version 10.0.8, Tree Star, OR, USA).

### IncuCyte Zoom imaging and image analysis

#### Image analysis of total image fluorescence intensity

Time-lapse fluorescence intensity data from Neuro-2a (HSE:EGFP) cells were acquired using an IncuCyte Live Cell Analysis System. Phase contrast and fluorescent images were acquired at 2 h intervals with the 10× or 20× objective. The fluorescence intensity of mCherry and stress-inducible EGFP were quantified using the basic analyser algorithm (Table 1) from a minimum of 9 images per well at each time point. Spectral overlap from the mCherry (3%) channel was removed from the EGFP channel in these images.

**Table 1.**
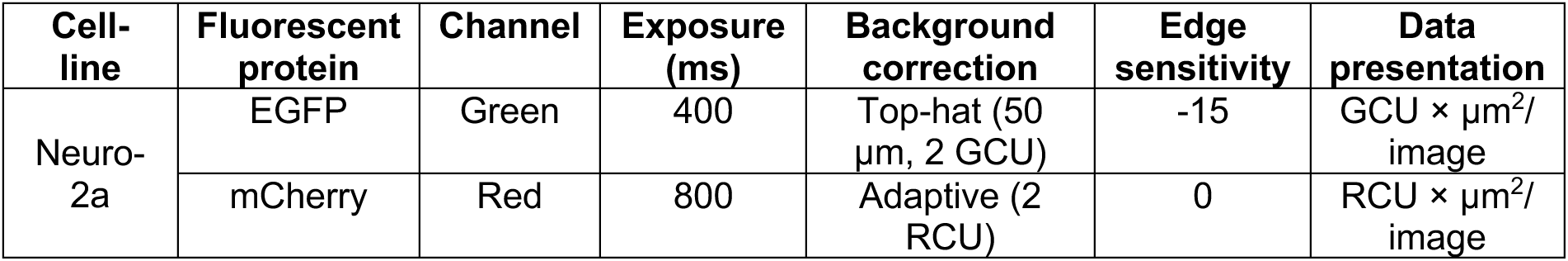
Cell mask parameters for the analysis of relative fluorescence intensities of EGFP and mCherry using the IncuCyte Zoom basic analyser.

| Cell-line | Fluorescent protein | Channel | Exposure (ms) | Background correction | Edge sensitivity | Data presentation |
| --- | --- | --- | --- | --- | --- | --- |
| Neuro-2a | EGFP | Green | 400 | Top-hat (50 μm, 2 GCU) | -15 | GCU × μm<sup>2</sup>/image |
| | mCherry | Red | 800 | Adaptive (2 RCU) | 0 | RCU × μm<sup>2</sup>/image |

The mean EGFP relative fluorescence intensity (RFU) was normalised by dividing the EGFP RFU by the mCherry RFU at each time point to account for relative changes in cell density over time (equation 1). The normalised EGFP data is presented as the mean fold change (Δ) in the EGFP/ mCherry ratio (± S.E.M.) of three independent repeats as described by equation 2, where *EGFP_Tx_* represents the EGFP RFU at any time and *EGFPT0* represents the EGFP RFU at 0 h.

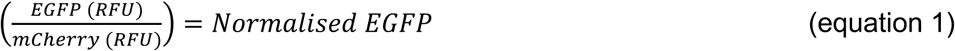

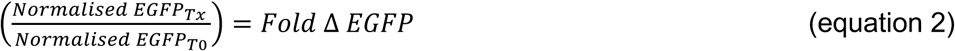

#### Image analysis of single cell fluorescent intensities

Neuro-2a (HSE:EGFP) were either left untransfected or transfected to express Htt^72Q^ or Fluc^DM^ in an 8-well Ibidi chamber, and imaged on a Leica SP5 confocal microscope at 37°C under 5% CO_2_/ 95% air. High-resolution images were captured using a 63 *×* water immersion objective and widefield images were captured using a 40 *×* air objective (see Supplementary Figure 4 for excitation and emission collection windows). Each well was imaged at 8 regions of interest at 1 h intervals for up to 60 h. The images obtained were analysed using CellProfiler 2.2.0 (Carpenter, Jones et al. 2006, Kamentsky, Jones et al. 2011, McQuin, Goodman et al. 2018). The mask parameters used for the identification of “cells” and “inclusions” are outlined in Table 2. These parameters were optimised to separate cell clumps and identify individual cells and inclusions. Using these parameters, the following custom-made sequence of image processing events was used to analyse all images in a non-biased manner; (i) all images were converted to greyscale, (ii) “cells” were identified as primary objects (Table 2), (iii) “inclusions” were identified as primary objects (Table 2), (iv) the region of the cell cytoplasm excluding the inclusions was defined as “cells – inclusions” tertiary objects, (v) the Cerulean fluorescence intensities of “cells” and EGFP fluorescence intensities of the regions defined as “cells – inclusions” was measured. The tertiary objects identified, “cells – inclusions”, were applied to eliminate the spectral overlap of the Cerulean fluorescence signal at the site of inclusions into the EGFP fluorescence signal. In this way EGFP fluorescence was measured from an area of the cell that did not contain inclusion bodies (i.e. “cells – inclusions”).

**Table 2.** Mask parameters for the analysis of relative fluorescence intensities of individual cells in confocal imaging experiments using Cell Profiler.

| Object identified | Input image | Diameter (pixel units) | Threshold strategy | Thresholding method | Threshold boundaries | Distinguish clumps | Dividing lines in clumps |
| --- | --- | --- | --- | --- | --- | --- | --- |
| Nuclei | Hoechst | 20-100 | Automatic | - | - | Intensity | Shape |
| Cells | mCherry | 20-65 | Global | Otsu | 0.01-1.0 | Intensity | Propagate |
|  | EGFP | - | Global | Otsu | 0.0-1.0 | - | - |
| Inclusions | Cerulean | 15-40 | Global | Otsu | 0.01-1.0 | Intensity | Shape |

Bivariate plots of the Cerulean and EGFP fluorescence intensities derived from regions defined as “cells – inclusions” demonstrated a strong correlation suggesting that the Cerulean signal was still contributing to the EGFP signal (Supplementary Figure 4c). This spectral overlap was calculated to be 13% based on EGFP- and Cerulean-only controls. Therefore, spectral compensation was performed on the EGFP data according to equation 3.

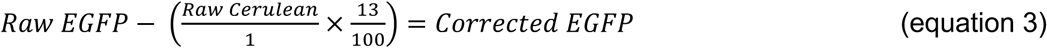

Lastly, thresholding was used to count the number of Cerulean^+ve^ (or transfected) cells or EGFP^+ve^ (or HSR^+ve^) cells at each time point. These thresholds were determined from the EGFP RFU and Cerulean RFU of cells in the untransfected and untreated samples (in this case 65 RFU and 15 RFU, respectively).

### Flow cytometric analysis

#### Analysis of whole cells

Flow cytometry was performed using an LSR Fortessa X-20 cell analyser equipped with 405 nm, 488 nm, 561 nm and 640 nm lasers (BD Biosciences, CA, USA). A minimum of 20,000 events per sample were collected at a high flow rate. Forward scatter was collected using a linear scale and side scatter in a log scale. Fluorescent emissions were collected as area (log scale), pulse height (log scale), and pulse width (linear scale) for each channel. For Cerulean fluorescence, data was collected with the 405 nm laser and 450/50 nm filter, EGFP was collected with the 488 nm laser and 525/50 nm filter, mCherry was collected with the 561 nm laser and 586/15 nm filter, and RedDot 1 was collected with the 640 nm laser and 670/30 nm filter. Spectral compensation, gating and data analysis of events acquired by flow cytometry was performed using Flow Jo software (Tree Star).

#### Flow cytometric analysis of inclusions and trafficking (FloIT)

Cells to be analysed were grown and transfected in 24-well plates. Cells were washed twice with PBS (pH 7.4) 48 h post-transfection, harvested mechanically by aspiration on ice, and resuspended in 500 µL ice-cold PBS for analysis of intact cells or cell lysates by flow cytometry. An aliquot of the cell suspension (150 µL) was taken and the transfection efficiency determined using untransfected cells as a negative control sample. The remaining 350 µL of cell suspension was lysed as described previously (Whiten, San Gil et al. 2016) in PBS containing a final concentration of 0.5% (v/v) TritonX-100 and 1 × Halt protease and phosphatase inhibitors. Except in control samples used to set gates, RedDot1 (Biotium, CA, USA) was diluted 1:1000 into lysis buffer prior to adding to cells. After 2 min incubation at room temperature to lyse cells, the lysate was analysed by flow cytometry measuring forward and side scatter, together with RedDot1 fluorescence. Analysis of all events was performed using Flow Jo (Tree Star). The number of inclusions in each sample was normalised to the number of transfected nuclei the same sample and the transfection efficiency was determined from the whole-cell data.

### Statistics

Results shown are the mean ± S.E.M. of three independent experiments unless otherwise indicated. Evaluation of statistical differences between the means of groups was determined by a one-way analysis of variance (ANOVA) or two-way ANOVA for multiple comparisons. The F-statistic from the ANOVA test and its associated degrees of freedom (between groups and within groups, respectively) are reported in parentheses. The *P*-value from the ANOVA test is also stated. Post hoc testing for differences between means was done using Dunnett’s when comparing means to the mean of the control, Tukey’s for multiple comparisons of the means, or Bonferroni’s for comparing the differences in the means between two samples over time, using GraphPad Prism 5 (GraphPad Software, Inc., CA, USA) and as described in the appropriate figure legends. For data showing the fold change in EGFP expression over time in Neuro-2a (HSE:EGFP) cells, a non-linear fit was applied [log(agonist) vs. response (Variable slope)] to the data. The time taken to reach half maximal EGFP intensity was determined by using the logEC50 value as a measure of the kinetics of the HSR.

## Results

### Generation and validation of an HSR reporter cell line

To investigate the activation of the HSR, stable Neuro-2a cell lines were generated in which expression of a fluorescent protein (EGFP) was used to report on HSR induction. These cells also constitutively express mCherry to account for changes in cell number or general transcription rates over time with different treatment paradigms. To validate that this stable cell line reports on HSR induction, the cells were first treated with known inducers of the HSR, namely CdCl_2_, heat shock, and celastrol (a compound identified in a drug screen as a potent inducer of the HSR) (Heemskerk, Tobin et al. 2002, Westerheide, Bosman et al. 2004). The fluorescent reporter, EGFP, has a half-life of 27 h (Corish and Tyler-Smith 1999), therefore these experiments were performed over a time-frame that would facilitate the accumulation of EGFP after HSR induction in order to assess the magnitude of the HSR. Concentration-response experiments were conducted to determine the concentration of CdCl_2_ (0-33 µM) and celastrol (0-1 µM) that induces a maximal HSR (Supplementary Figure 1). Based on these results, subsequent experiments used 10 µM CdCl_2_ and 0.75 µM celastrol over a 24 h time-frame, which were the lowest concentrations that induced a maximal HSR in Neuro-2a (HSE:EGFP).

Following treatment with 10 µM CdCl_2_, heat shock (42°C for 2 h), or 0.75 µM celastrol, there was a time-dependent increase in EGFP fluorescence intensity in Neuro-2a (HSE:EGFP) cells (Figure 1a and b). Fluorescence was first detected 2-4 h following heat shock treatment of Neuro-2a (HSE:EGFP) cells (Figure 1b), and reached maximal response after 12 h. The time taken to induce the HSR and to reach maximum EGFP fluorescence varied between treatment types. The expression of EGFP was induced significantly faster after heat shock (5.1 ± 0.2 h) compared to treatment with CdCl_2_ or celastrol (11 ± 0.6 h and 8.3 ± 1.5 h, respectively; Figure 1c) [F (2, 6) = 28.28, *P* = 0.0009] as measured by the time taken to reach half maximal fluorescence. Treatment with each of these classical inducers of the HSR resulted in a significant increase in the magnitude of EGFP expression in Neuro-2a (HSE:EGFP) cells compared to no treatment [F (3,8) = 4.265, *P* = 0.0448] (Figure 1d). However, there were no differences observed between the magnitude of HSR induction between treatment type in Neuro-2a (HSE:EGFP) cells as determined by the fold change in fluorescence intensity at 24 h (Figure 1d). Beyond the 24 h time-frame of these experiments, cells treated with CdCl_2_ or celastrol died as a consequence of the toxicity of these treatments. Heat shocked cells continued to grow and divide during the recovery period; EGFP fluorescence intensity peaked at 10 h.

**Figure 1.**
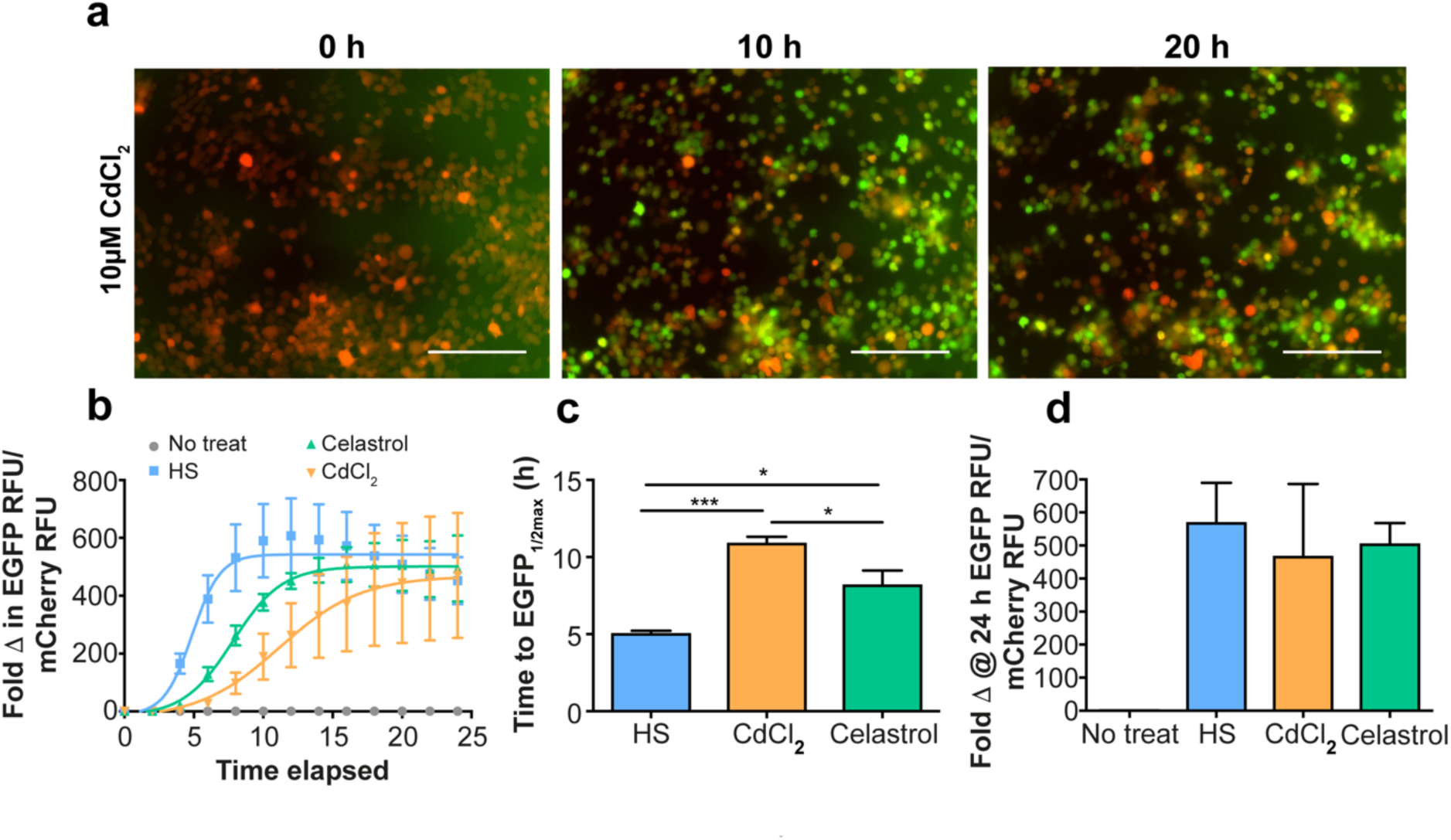
Neuro-2a (HSE:EGFP) cells enable specific and sensitive quantification of HSR induction. (a) Neuro-2a (HSE:EGFP) cells stably expressing mCherry were treated with 10 µM CdCl_2_ and imaged every 2 h to monitor EGFP expression. Representative overlay images of mCherry and EGFP fluorescence are shown after 0, 10 and 20 h of treatment. Scale bars = 200 μm. (b) The fold change in EGFP fluorescence intensity over time normalised to mCherry fluorescence intensity to account for changes in cell density during the experiment. The HSR was induced in these cells by heat shock (42°C for 2 h) or treatment with 0.75 µM celastrol or 10 µM CdCl_2_. (c) The kinetics of HSR induction as determined by the time taken to reach half maximal EGFP fluorescence. (d) The magnitude of HSR induction as determined by EGFP fluorescence after 24 h of treatment with each stress. Data shown are the mean ± S.E.M. of three independent repeats. Differences between the means were assessed using a one-way ANOVA followed by Tukey’s post-hoc test, where *P* < 0.05 (*), *P* < 0.01(**), and *P* < 0.001 (***).

### Extracellular protein aggregates do not induce an HSR

There is increasing evidence that disease-associated protein aggregation propagates through the CNS from the site of onset in a prion-like mechanism. This process likely involves the release of misfolded and aggregated protein from neurons into the extracellular space and their subsequent uptake by surrounding cells (Jucker and Walker 2013, Zeineddine, Pundavela et al. 2015, Zeineddine and Yerbury 2015). We therefore sought to determine whether cells respond to the extracellular application of aggregated protein by inducing an HSR. To test this, recombinant human α-synuclein and SOD1^G93A^ were aggregated *in vitro*. There was a significant increase in ThT fluorescence of the aggregated α-synuclein sample compared to monomeric α-synuclein and buffer alone, indicative of an increase in β-sheet structure with aggregation (Figure 2a). Transmission electron microscopy (TEM) indicated that the α-synuclein had formed long, mature fibrils > 2 µm in length (Figure 2a inset). The formation of SOD1^G93A^ aggregates was monitored by an *in situ* ThT binding assay; there was a time-dependent increase in ThT fluorescence relative to buffer alone (Figure 2b). The SOD1^G93A^ aggregates formed in these assays had an irregular amorphous structure and were < 80 nm in length (Figure 2b inset). Together, the ThT and TEM data indicate that the SOD1^G93A^ aggregates formed in this study contain an underlying β-sheet structure, but did not assemble into highly-ordered fibrils.

**Figure 2.**
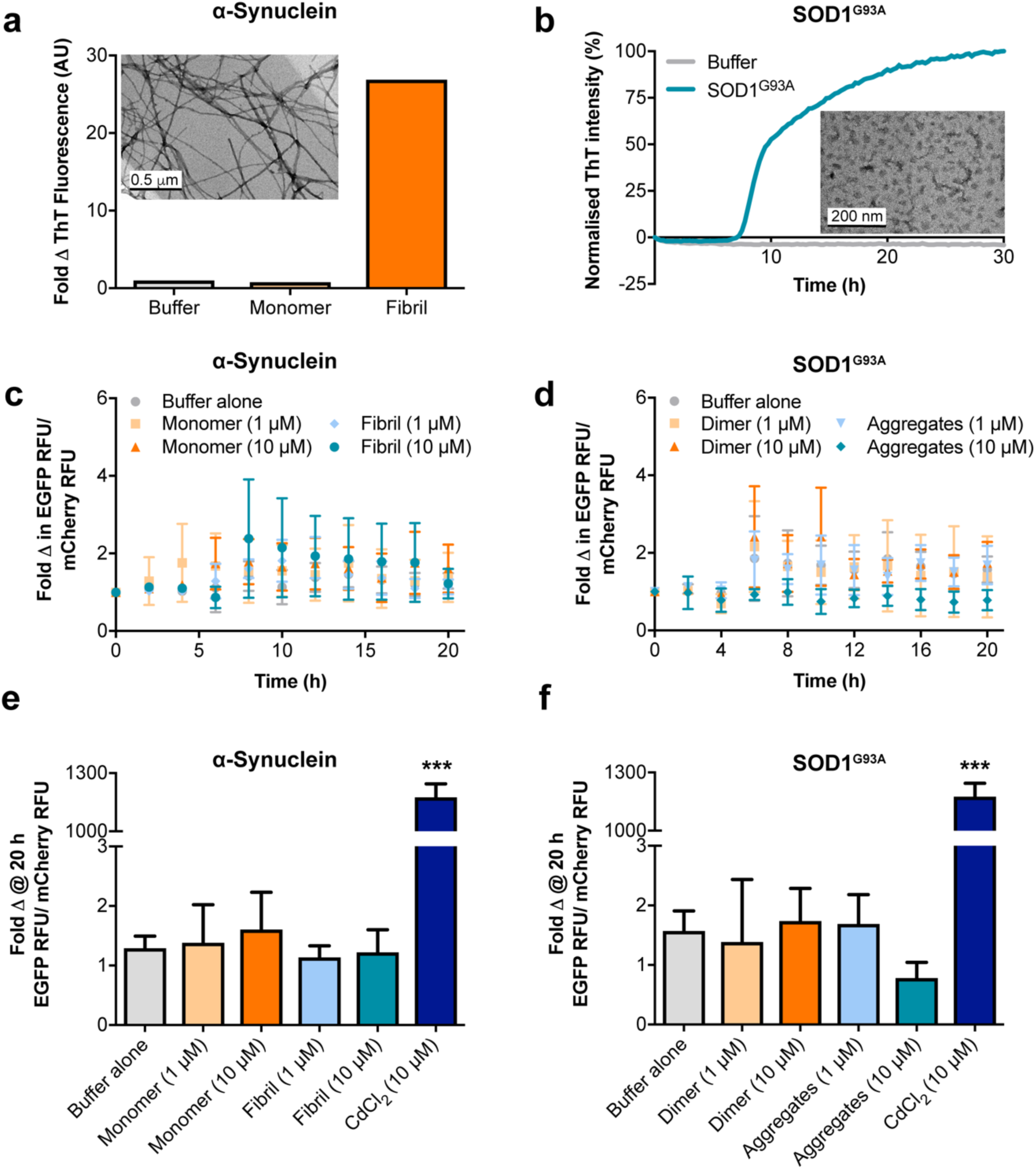
Disease-associated protein aggregates do not induce an HSR in Neuro-2a (HSE:EGFP) when applied extracellularly. The formation of (a) α-synuclein fibrils, and (b) SOD1^G93A^ aggregates was confirmed by ThT fluorescence and TEM (insets) (scale bar = 500 and 200 nm, respectively). (c-d) Monitoring induction of the HSR using live-cell imaging. Time course of EGFP RFU normalised to mCherry RFU of Neuro-2a (HSE:EGFP) cells treated with (c) monomeric or fibrillar α-synuclein, or (d) dimeric or aggregated SOD1^G93A^. (e-f) The magnitude of HSR activation as determined by the EGFP RFU/ mCherry RFU 20 h after application of the protein aggregates or CdCl_2_. Data shown are the mean ± S.E.M. of at least three independent repeats. Differences between the means were assessed using a one-way ANOVA followed by Dunnet’s post-hoc test (comparing to buffer alone control), where *P* < 0.001 (***).

Non-aggregated and aggregated forms of α-synuclein and SOD1^G93A^ were applied to Neuro-2a (HSE:EGFP) and the cells monitored for HSR induction by time-lapse live-cell imaging (Figure 2c-f). There was no significant difference in EGFP fluorescence in cells treated for 20 h with either 1 µM or 10 µM of non-aggregated or aggregated α-synuclein or SOD1^G93A^ (Figure 2e-f), and this remained the case even up to 72 h after treatment (data not shown). In contrast, the positive control CdCl_2_ treatment induced an HSR in these cells (Figure 2e-f). Thus, these extracellular aggregates of α-synuclein and SOD1^G93A^ associated with neurodegenerative disease were not sufficient to induce an HSR in these cells.

### Intracellular aggregation of pathogenic proteins is a poor inducer of the HSR

We next investigated whether the aggregation of proteins into inclusions in cells leads to the induction of an HSR in Neuro-2a (HSE:EGFP) cells. To examine this, a suite of constructs was generated for the expression of Cerulean-tagged SOD1^WT^, SOD1^G93A^, Htt^25Q^, Htt^72Q^, Fluc^WT^, or Fluc^DM^. These proteins were selected because they represent a mix of (i) pathogenic (SOD1 and Htt) and non-pathogenic (Fluc) aggregation-prone proteins and their WT isoforms, (ii) JUNQ-, iPOD-, and “other” inclusion forming proteins (SOD1, Htt, Fluc, respectively), and (iii) amyloidogenic (SOD1 and Htt) and amorphous (Fluc) proteins.

The propensity of each of these Cerulean-tagged proteins to aggregate was assessed by FloIT, a flow cytometric method for the quantification of inclusions (Whiten, San Gil et al. 2016). The addition of RedDot1 to cell lysates enabled nuclei to be identified and enumerated (Figure 3a-b). Lysates from cells transfected to express SOD1^WT^, which does not readily form inclusions (McAlary, Aquilina et al. 2016, Whiten, San Gil et al. 2016), were used as a negative control for inclusion formation (Figure 3c). Cerulean-tagged SOD1^G93A^ formed inclusions, which could be quantified (Figure 3d-e). The total number of inclusions formed by each over-expressed protein varied and was highest for Fluc^WT^ and Fluc^DM^. Differences between the means were determined using a one-way ANOVA followed by Tukey’s post-hoc test and there were no statistically significant differences in the number of aggregates quantified in each sample.

**Figure 3.**
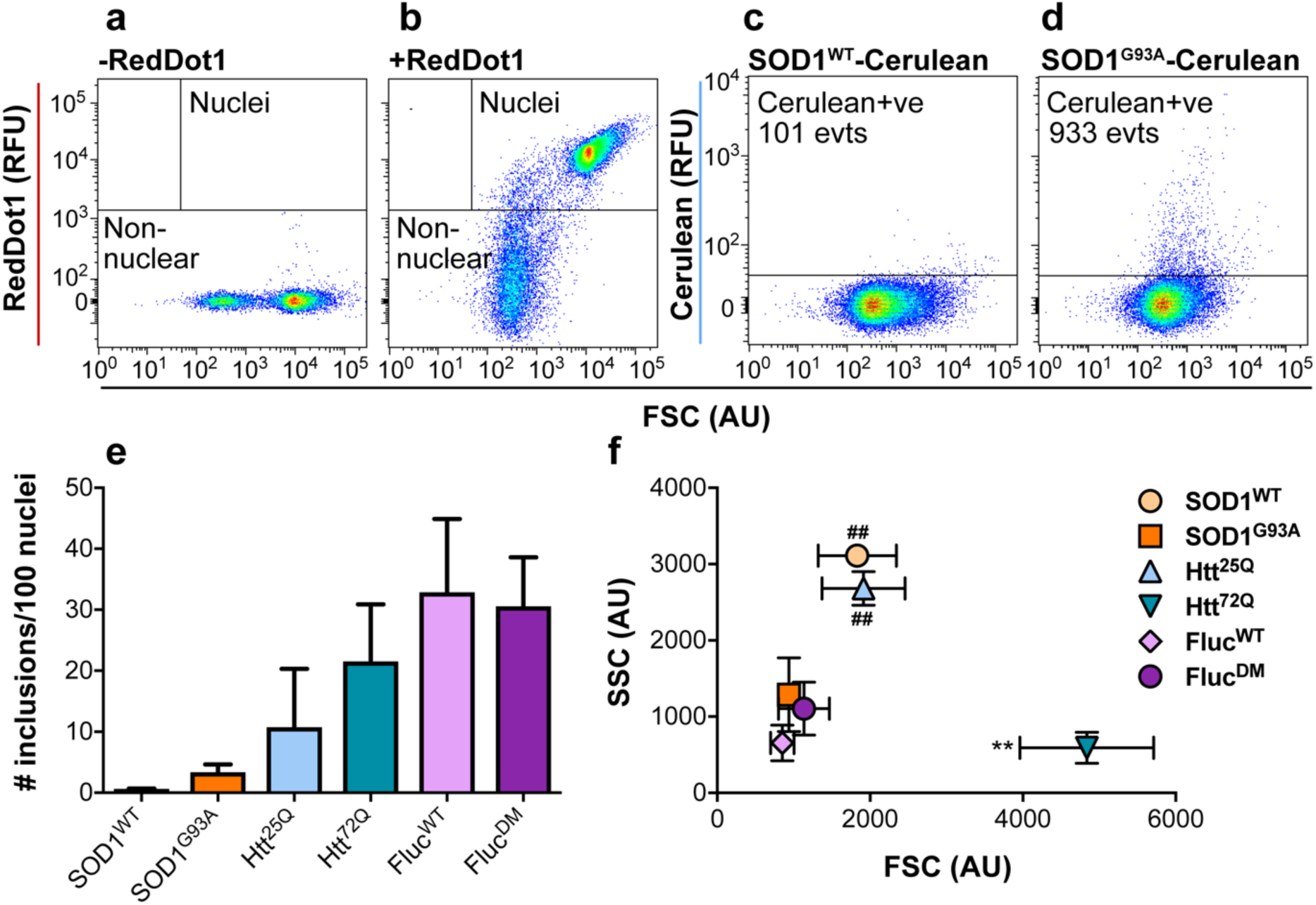
Flow cytometric analysis of cell lysates (FloIT) demonstrates that different pathogenic and non-pathogenic proteins form different quantities and sizes of inclusions in Neuro-2a (HSE:EGFP) cells. Neuro-2a (HSE:EGFP) cells were transfected to express Cerulean-tagged SOD1^WT^, SOD1^G93A^, Htt^25Q^, Htt^72Q^, Fluc^WT^, or Fluc^DM^ and, 48 h post-transfection, cells were lysed and analysed by FloIT. (a)-(b) Plots of forward scatter (FSC; size) and RedDot1 nuclear dye fluorescence used to enumerate the nuclei in the cell lysates. (a) Cells lysed in the absence of RedDot1 were used to set square gates to capture RedDot1^+ve^ events and RedDot^-ve^ non-nuclear events. (b) Cells lysed in the presence of RedDot1 are shown as a representative plot. (c)-(d) Plots of FSC and Cerulean RFU. Representative plots show events acquired from Neuro-2a (HSE:EGFP) cells transfected to express SOD1^WT^-Cerulean and SOD1^G93A^-Cerulean. The number of Cerulean^+ve^ events are denoted in the gate. (e) The number of Cerulean^+ve^ inclusion bodies quantified by FloIT, normalised to the number of transfected nuclei (total number of nuclei divided by the transfection efficiency of whole cells). (f) Bivariate plot of the FSC (size) and side scatter (SSC; granularity) of the Cerulean^+ve^ inclusions identified by FloIT. Data shown are the mean ± S.E.M. of three independent repeats, some error bars were small and can’t be seen on the plot. Differences between the means were determined using a one-way ANOVA followed by Tukey’s post-hoc test, there were no statistically significant differences in (e) and in (f) FSC, *P* < 0.01 (**) and SSC, *P* < 0.01 (##).

One factor that could possibly influence HSR induction is the size and granularity of the inclusions formed by each protein, since this would be an indirect measure of the surface area available to interact with cytoplasmic proteins. FloIT analysis demonstrated that the SOD1^WT^ and Htt^25Q^ aggregates detected had a significantly greater granularity (SSC) compared to the other proteins tested (*P* <0.01). In addition, the relative size of the inclusions formed by Htt^72Q^ were significantly larger (based on the forward scatter, FSC, signal, Figure 3f) compared to inclusions formed by Fluc^DM^ and SOD1^G93A^ [F (5, 8) = 8.027, *P* = 0.0056].

Cells over-expressing these Cerulean-tagged proteins were also analysed by flow cytometry to assess whether inclusion formation by aggregation-prone proteins induced an HSR in Neuro-2a (HSE:EGFP) cells (Supplementary Figure 2). Untransfected cells were used as an EGFP^-ve^ control (Figure 4a) and cells treated with celastrol (0.75 µM/ 24 h; Figure 4b) used as an EGFP^+ve^ control in these experiments. Representative flow cytometric data of the proportion of EGFP^+ve^ cells is shown for cells expressing Fluc^DM^ in Figure 4c. Expression of the two pathogenic proteins, SOD1^G93A^ or Htt^72Q^, resulted in 1.6 ± 0.4% and 4.0 ± 1.1% of the transfected cells becoming EGFP^+ve^, respectively (Figure 4d and Supplementary Figure 3). However, this was not significantly different than the proportion of EGFP^+ve^ cells expressing non-aggregation prone isoforms of these proteins, i.e. SOD1^WT^ or Htt^25Q^. In contrast, the expression of Fluc^DM^ resulted in HSR induction in 12.3 ± 1.3% of transfected cells, which was significantly greater than HSR induction in cells expressing the aggregation-prone disease-related proteins (Figure 4c-d).

**Figure 4.**
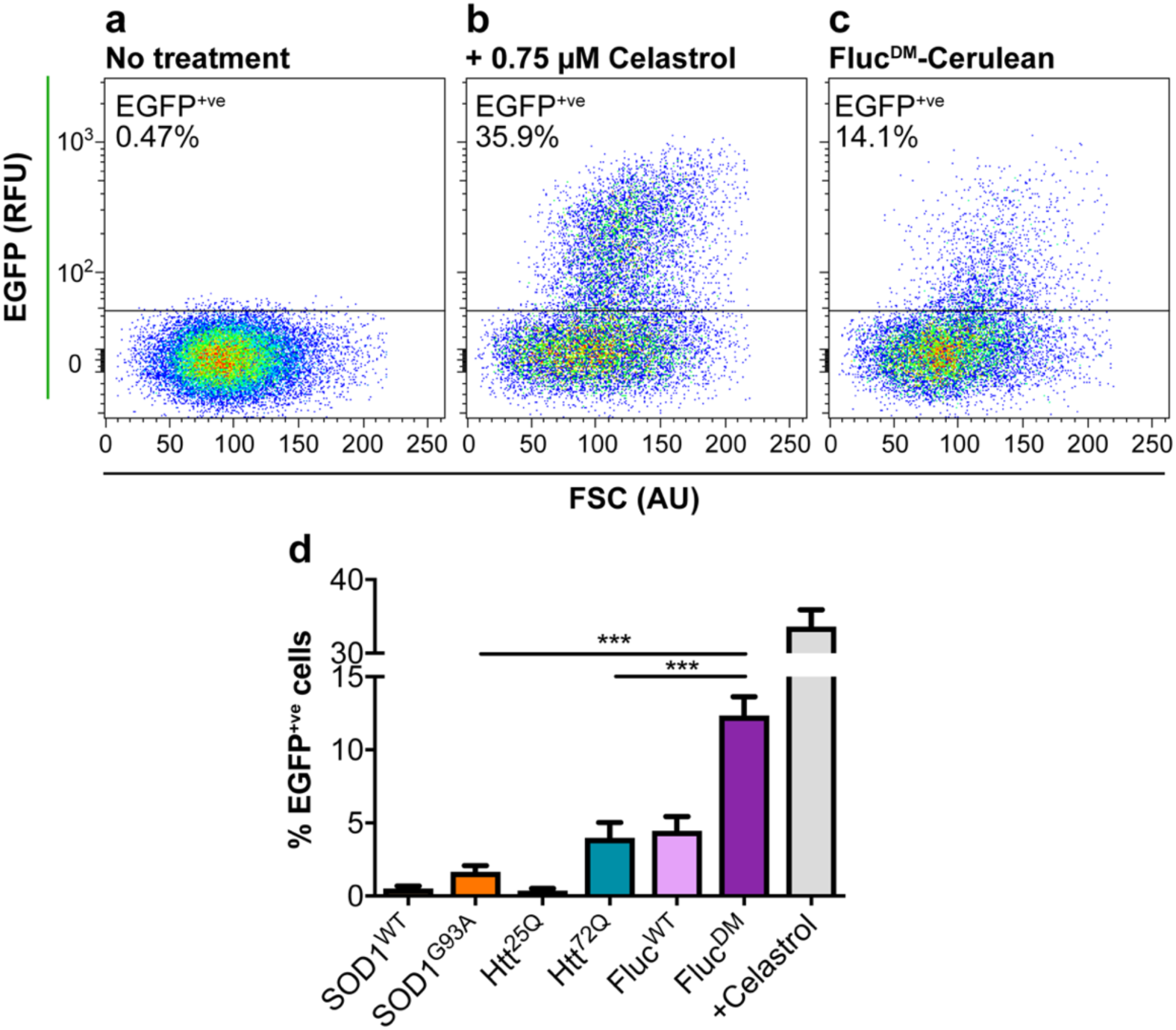
Over-expression of Fluc^DM^, but not disease-associated proteins, induces the HSR in Neuro-2a (HSE:EGFP) cells. Neuro-2a (HSE:EGFP) cells were transfected to express Cerulean tagged SOD1^WT^, SOD1^G93A^, Htt^25Q^, Htt^72Q^, Fluc^WT^ or Fluc^DM^ proteins. After 48 h incubation, cells were harvested for analysis by flow cytometry. Cellular debris, cell clumps, doublet events and untransfected cells were excluded from the analysis as described in Supplementary Figure 2. (a)-(c) Representative plots of FSC and EGFP fluorescence of (a) untransfected and untreated cells, (b) cells treated with 0.75 µM celastrol (an inducer of the HSR), and (c) cells transfected to express Fluc^DM^-Cerulean. (d) The percent of transfected cells expressing Cerulean-tagged WT or mutant proteins which were EGFP^+ve^. Data shown are the mean ± S.E.M. of three independent repeats. Statistically significant differences between the means were determined by one-way ANOVA and Tukey’s multiple comparisons test, where *P* < 0.001 (***).

Given that the analysis of the entire cell population demonstrated that pathogenic SOD1 and Htt did not elicit a significant HSR compared to their respective WT proteins, we next sought to determine whether the expression level of an aggregation-prone protein affects the induction of the HSR in these cells. To do so, events from the flow cytometric analyses were binned according to the levels of Cerulean expression (5000 RFU per bin, where bin 1 contains cells with the lowest expression and bin 10 the highest level of expression; Figure 5a). For cells in each of these bins the proportion of cells in which an HSR was induced was determined. Representative bivariate plots of EGFP and FSC of cells transfected to express Htt^72Q^ in Cerulean bins 5 and 10 are shown in Figure 5b. By binning the flow cytometry data in this way two factors can be assessed: (i) the effect of high and low protein concentration on induction of the HSR, and (ii) the induction of the HSR in cells expressing WT and mutant aggregation-prone proteins (Figure 5c-e).

**Figure 5.**
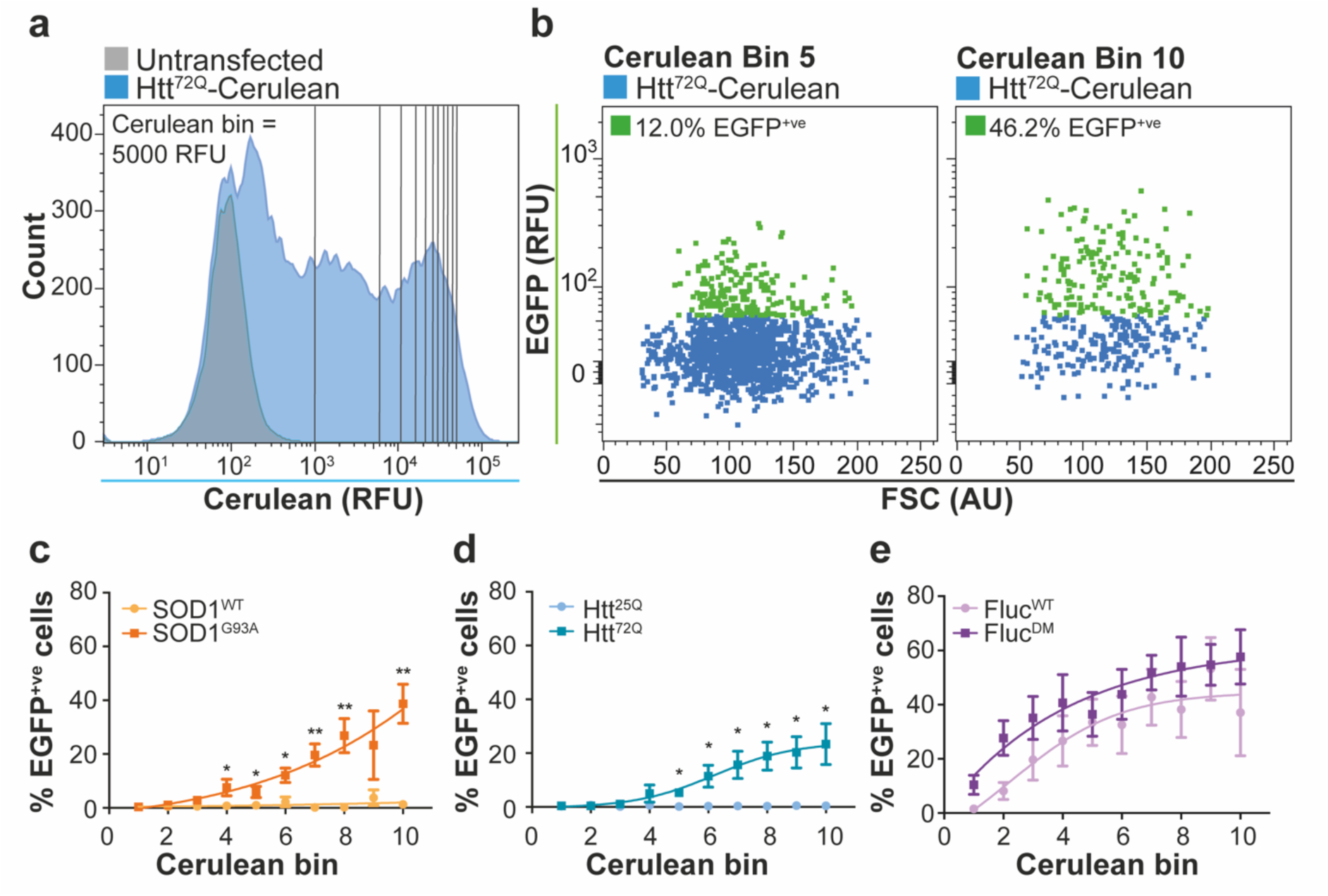
High expression levels of aggregation-prone proteins correlate with an increase in the proportion of cells with an activated HSR. (a) Overlay histograms of untransfected (*grey*) and Htt^72Q^-Cerulean (*blue*) transfected Neuro-2a (HSE:EGFP) with 10 Cerulean bins corresponding to 5000 AFU. (b) Representative plots of FSC and EGFP fluorescence from cells transfected to express Htt^72Q^-Cerulean (bin 5, *left*; bin 10, *right*). The EGFP^+ve^ cells in the indicated Cerulean bin (*green*) are highlighted. (c)-(e) The percent of EGFP^+ve^ cells in each Cerulean bin for cells expressing (c) SOD1^WT^ or SOD1^G93A^, (d) Htt^25Q^ or Htt^72Q^, or (e) Fluc^WT^ or Fluc^DM^. Data shown are the mean ± S.E.M. of three independent repeats. Differences between the means were determined using a two-way ANOVA followed by Bonferroni’s post hoc test, where *P* < 0.05 (*) and *P* < 0.01 (**).

In SOD1^G93A^-Cerulean expressing cells, as the amount of SOD1^G93A^ expressed by cells increased so did the proportion of cells in which an HSR had been induced (0.4 ± 0.2% of cells were EGFP^+ve^ in bin 1 compared to 38.7 ± 7.3% of cells in bin 10; Figure 5c). This same trend was observed for cells expressing Htt^72Q^, Fluc^WT^ and Fluc^DM^, *i.e.* cells expressing the highest amounts of these proteins (bin 10) had the highest proportion of EGFP^+ve^ cells (23.4 ± 7.6%, 37.1 ± 15.9%, and 57.7 ± 10%, respectively; Figure 5d-e). In contrast, over-expression of non-aggregation prone isoforms of SOD1^WT^ and Htt^25Q^ did not induce an HSR in any cerulean bin demonstrating that it is not simply protein over-expression that induces an HSR. The HSR was most sensitive to increasing concentrations of Fluc^DM^ and Fluc^WT^, for example in Cerulean bins 1 and 2, respectively, an EGFP^+ve^ population was already evident. Comparatively, there is a significant increase in the proportion of EGFP^+ve^ cells in bin 5 and bin 4 for Htt^72Q^ and SOD1^G93A^, respectively, compared to WT controls. Together, these data show that an HSR is activated at lower relative levels of expression of Fluc^WT^ and Fluc^DM^ compared to Htt^72Q^ and SOD^G93A^.

### Formation of inclusions precedes the detection of HSR induction

Finally, we sought investigate whether HSR activation occurs before or after inclusion formation in cells to determine whether the HSR induction we observe is in response to protein aggregation. We used Neuro-2a (HSE:EGFP) cells over-expressing Htt^72Q^ or Fluc^DM^ for these experiments as both readily form inclusions yet have a differential capacity to induce an HSR (Figure 3e and Figure 4d). This is exemplified by the proportion of EGFP^+ve^ events in cells expressing Htt^72Q^ (4.0 ± 1.1%) compared to Fluc^DM^ (12.3 ± 1.3%; Figure 4d). Live-cell time-lapse confocal imaging of Neuro-2a (HSE:EGFP) cells facilitated the simultaneous tracking of Htt^72Q^ or Fluc^DM^ expression, inclusion formation, and HSR induction in single cells (Figure 6).

**Figure 6.**
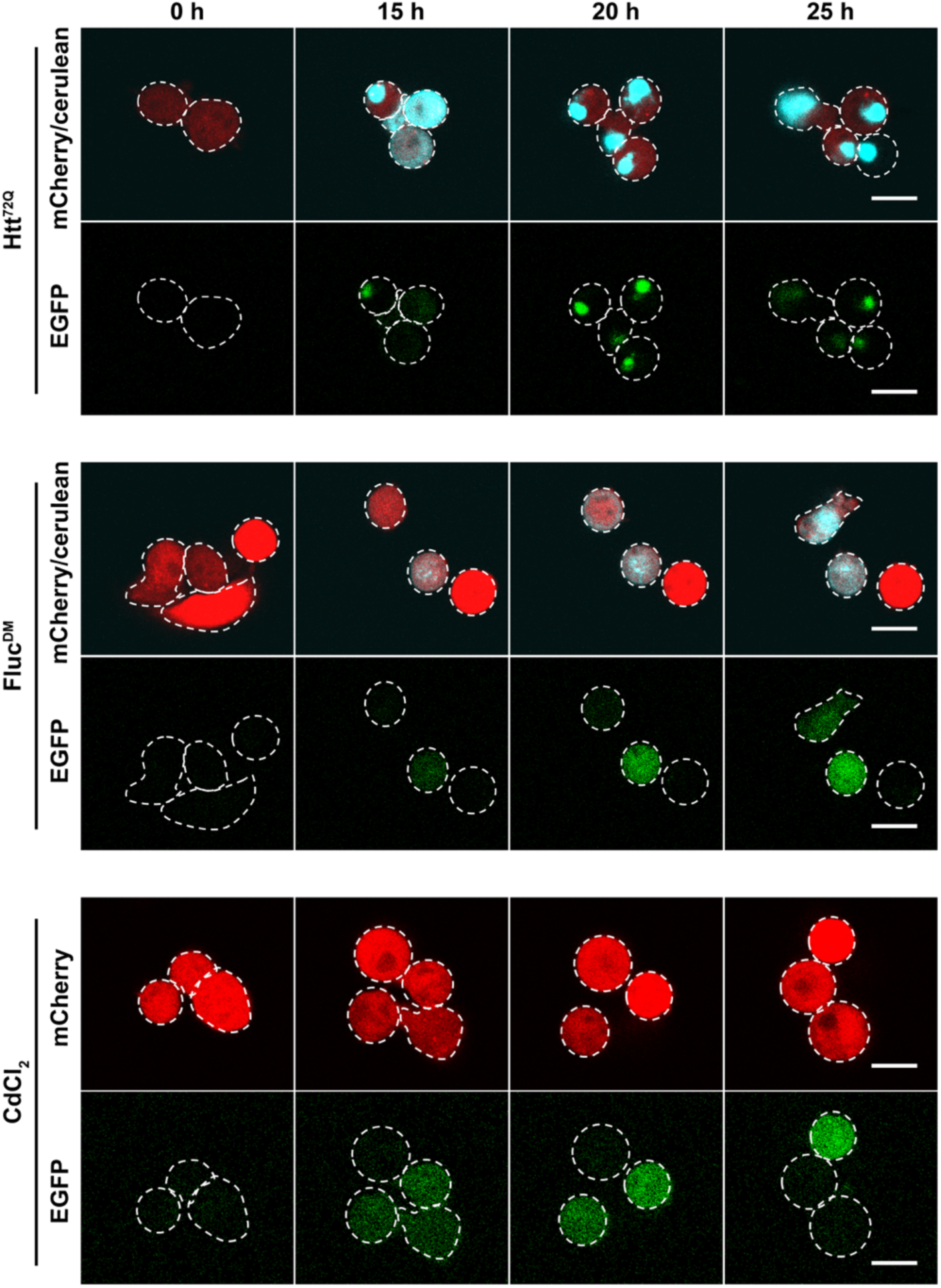
The induction of the HSR and inclusion body formation captured by live cell imaging. Neuro-2a (HSE:EGFP) cells were transfected to express Htt^72Q^ (top panel) or Fluc^DM^ (middle panel), or treated with 10 µM CdCl_2_ (bottom panel) and imaged every hour by confocal microscopy. Representative confocal images are shown after 0, 15, 20, and 25 h of treatment. Each image is overlayed by the cell outlines (white dotted line) as defined by the mCherry signal at each time point. Punctate EGFP signal represents spectral overlap from the Cerulean signal and diffuse EGFP signal represents the activation of the HSR. Scale bar = 20 µm. Images are representative of three independent experiments.

There was no detectable HSR induction observed at any time point following transfection of cells with Htt^72Q^ (Figure 6). However, in cells expressing Fluc^DM^, there was an increase in EGFP expression, indicative of HSR induction, in 2 of the 3 representative cells depicted in Figure 6 after 15 h of incubation (Figure 6).

We used a non-biased image quantification strategy to analyse wide-field of view images of treated or transfected cells over time to assess the proportion of HSR-positive cells. Due to the broad fluorescence emission spectrum of the Cerulean protein, spectral overlap was observed between the channels used to detect EGFP and Cerulean fluorescence, particularly in regions containing inclusions (Supplementary Figure 4a). To minimise spectral overlap, narrow emission windows for detection of Cerulean and EGFP fluorescence were used when imaging (462-492 nm and 506-563 nm, respectively; Supplementary Figure 4a). Despite this, an apparent EGFP signal was observed in areas containing inclusions comprised of Cerulean-tagged proteins (areas of intense fluorescence; Supplementary Figure 4a inset; white arrowheads), as a result of Cerulean fluorescence being detected in the EGFP emission window. Thus, to accurately measure the level of EGFP over-time, “cells” and “inclusions” were identified based on their mCherry and Cerulean fluorescence, respectively, and EGFP fluorescence intensity was measured from an area in the cell defined as “cells – inclusions” (Supplementary Figure 4b).

Images from these live-cell imaging experiments were subjected to the analyses outlined in Supplementary Figure 4, to track the expression of Cerulean-tagged proteins, formation of inclusions, and induction of the HSR over time. Treatment of Neuro-2a (HSE:EGFP) with 10 µM CdCl_2_ (the positive control) resulted in a time-dependent increase in the proportion of cells that were EGFP^+ve^ compared to untreated cells (Figure 7a). Treatment with CdCl_2_ significantly increased the proportion of EGFP^+ve^ cells compared to no treatment [F (1, 130) = 369.7, *P* < 0.0001], and this reached statistical significance 12 h after treatment. Likewise, there was a time-dependent increase in the proportion of cells with an active HSR in samples transfected to express Fluc^DM^ (Figure 7b). In contrast, there was no or low effect of Htt^72Q^ expression on the proportion of EGFP^+ve^ cells over time, which indicates the relative lack of HSR induction in these cells (Figure 7b). There was a significantly greater proportion of EGFP^+ve^ cells in samples expressing Fluc^DM^ compared to Htt^72Q^ [F (1, 144) = 33.43, *P* < 0.0001]. Post-hoc analysis using Bonferroni’s test showed that this difference was statistically significant 21 h following transfection. Therefore, Fluc^DM^ and Htt^72Q^ have differential capacities to induce the HSR.

**Figure 7.**
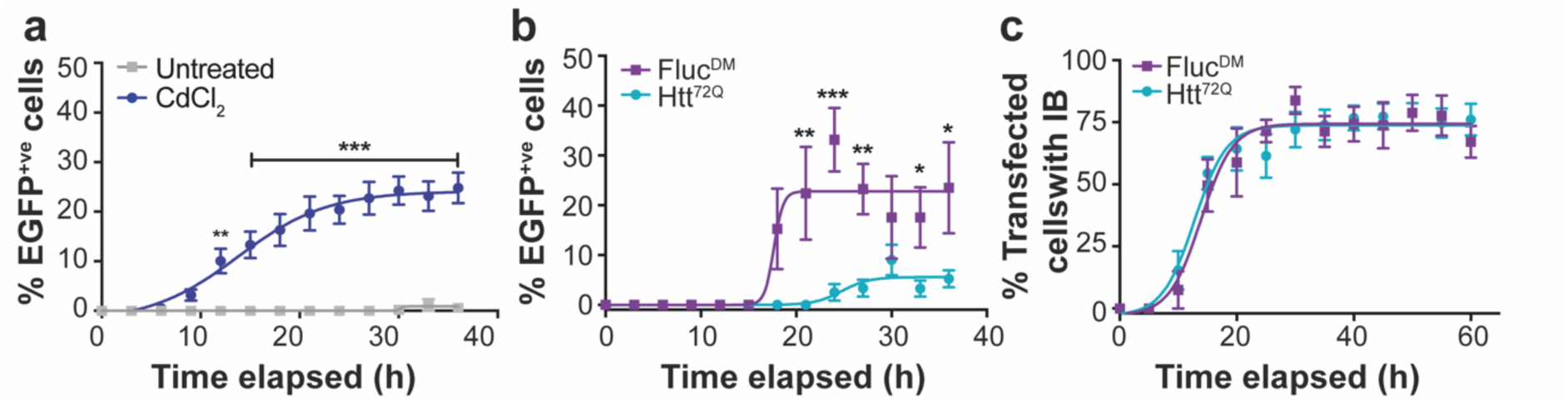
The induction of the HSR in cells expressing Fluc^DM^ or Htt^72Q^ is not dictated by the rate of aggregate formation. (a) The proportion of EGFP^+ve^ cells over time after treatment (or not) with 10 µM CdCl_2_. (b) The proportion of EGFP^+ve^ cells over time after transfection to express Htt^72Q^ or Fluc^DM^. (c) The percent of transfected cells with inclusions over time. Data shown are the mean ± S.E.M. of single cell analyses of >300 transfected cells from 8 fields of view using a 40× dry objective. These findings are representative of three independent live-cell imaging experiments. Differences in the means were assessed using a two-way ANOVA followed by post-hoc analysis using Bonferroni’s test, where *P* < 0.05 (*), *P* < 0.01 (**), and *P* < 0.001 (***).

To determine whether this difference in HSR induction was associated with the aggregation rate of Fluc^DM^ compared to Htt^72Q^, the proportion of cells with Htt^72Q^ or Fluc^DM^ inclusions was determined over the time-course of the live-cell imaging experiment (Figure 7c). There was a significant time-dependent increase in the number of inclusions formed in Neuro-2a (HSE:EGFP) by both Htt^72Q^ and Fluc^DM^, with both samples reaching a plateau in the proportion of cells with inclusions after 20 h [F (12, 143) = 33.43, *P* < 0.0001]. There was no effect of the type of protein expressed on the mean proportion of cells with inclusions formed over the time-course of the experiment [F (1, 143) = 0.09, *P* = 0.7607]. Both Htt^72Q^ and Fluc^DM^ reached half maximal inclusion formation 12.8 ± 2.4 h and 13.9 ± 3.4 h following transfection, respectively, indicating that there was no significant difference in the rate of aggregation of either protein. In addition, the first Fluc^DM^ inclusions are detected 10 h after transfection, whereas the first HSR-positive cells are detected 18 h after transfection (*i.e.* 8 hours later). This time delay from inclusion formation to HSR induction in cells expressing Fluc^DM^ likely represents the time taken for HSF1 to activate, translocate to the nucleus, and drive transcription of the EGFP reporter to a level sufficient for detection by the confocal microscope. One of the limitations of time-lapse image analysis was the ability to detect inclusions of a range of sizes and fluorescence intensities. Consequently, it is likely that the reported maximum proportion of cells with inclusions (75% after 25 h for Fluc^DM^ and Htt^72Q^) is an underestimate. Visual assessments of images confirmed that most transfected cells contained inclusions 36 h after transfection.

## Discussion

The HSR is widely recognised as a first line of defence against protein aggregation. However, an explanation as to why Hsps fail to suppress the formation of protein aggregates in neurodegenerative diseases is lacking. Therefore, we investigated whether inclusions are capable of inducing an HSR in a cell-line of neuronal origin. In this work, we provide strong evidence that extracellular and intracellular disease-associated protein aggregates are poor inducers of the HSR. Namely, using Neuro-2a (HSE:EGFP) cells, we showed that the extracellular application of pathogenic aggregates (*α*-synuclein and SOD1^G93A^) does not induce an HSR. Over-expression of pathogenic proteins, Htt^72Q^ and SOD1^G93A^, induced an HSR in a small but not significant proportion of cells, whereas the expression of the non-disease related Fluc^DM^ induced an HSR in a significantly greater proportion of cells compared to SOD1^G93A^ and Htt^72Q^. Moreover, we established that there is a positive relationship between the concentration of aggregation-prone proteins and induction of the HSR in cells, indicating that relatively higher intracellular concentrations of aggregation-prone proteins correlate with HSR induction. Our study provides evidence that disease-associated protein aggregates have limited capacity to induce an HSR. Therapeutic strategies that activate the HSR in order to enhance levels of molecular chaperones, such as the Hsps, and restore proteostasis may therefore be of benefit in these disorders.

The cell-to-cell transfer of protein aggregates through the extracellular medium is thought to be one mechanism by which neurodegenerative diseases progress from a focal point of onset (Scheckel and Aguzzi 2018). Indeed, both SOD1 and *α*-synuclein have been shown to act in a prion-like manner in various cell and animal models of disease [reviewed recently in (Leak, Frosch et al. 2019, McAlary, Plotkin et al. 2019)]. In the present study we showed that the extracellular application of SOD1^G93A^ and α-synuclein aggregates to Neuro-2a (HSE:EGFP) cells did not result in an induction of the HSR up to 72 h post-treatment. Previous studies have shown that treatment of other cell-types in this way elicits other stress-responses, such as inflammatory pathways. For example, treatment of EOC.13 microglial-like cells with SOD1^G93A^ aggregates resulted in an upregulation of TNFα (Roberts, Zeineddine et al. 2013). The absence of an HSR in cells after treatment with protein aggregates suggests that these pathogenic proteins evade detection by the components of the HSR that activate this pathway. Thus, the failure of aggregated SOD1 and α-synuclein to activate the HSR may contribute to disease progression in neurodegenerative diseases by enabling the seeding of inclusion formation in neighbouring cells that take up aggregates. Future research could investigate whether the pharmacological induction of the HSR prevents inclusion formation after seeding with aggregates, since this may be a therapeutic strategy to stop the progression of neurodegenerative diseases in the CNS. Indeed, a recent clinical trial has shown that the HSR-inducing compound, arimoclomol, is safe and potentially effective in patients with rapidly progressing SOD1-associated ALS (Benatar, Wuu et al. 2018).

One possible explanation for the observed differences in HSR induction across the different proteins tested could be the intrinsic propensity of the proteins to form amorphous or amyloid inclusions, or alternatively iPOD or JUNQ formation. Here we showed that the intracellular expression of the pathogenic proteins, SOD1^G93A^ and Htt^72Q^, resulted in a low proportion of cells (2-4% of transfected cells) that had induced an HSR 48 h after transfection. This suggests that the HSR does not detect early protein misfolding and subsequent inclusion formation in the majority of SOD1^G93A^ or Htt^72Q^ expressing cells. In comparison, a significantly greater proportion of cells expressing Fluc^DM^ induced an HSR. Interestingly, *in vitro*, mutant SOD1, *α*-synuclein (Figure 2), and Htt (Scherzinger, Lurz et al. 1997) are *β*-sheet forming, amyloidogenic proteins, whereas Fluc forms amorphous aggregates (Gupta, Kasturi et al. 2011). Therefore, perhaps the off-folding pathway (*i.e.* amorphous or amyloid aggregation), rather than the sub-type of inclusion (*i.e.* iPOD, JUNQ, or other), can explain the observed differences in HSR induction across the different proteins tested. This could be further examined by analysing a broader range of proteins that aggregate via amorphous and amyloid pathways.

Previous research using a HEK293 fluorescent reporter cell line suggested that expression of a pathogenic form of Htt, namely Htt^91Q^, did not lead to detectable induction of the HSR, irrespective of the expression level or aggregation status of the protein in the cell (Bersuker, Hipp et al. 2013). Likewise, over-expression of artificial *β*-sheet forming proteins in HEK293T cells did not induce the expression of Hsp110, Hsp70 or Hsp27, markers of HSR induction (Olzscha, Schermann et al. 2011). Moreover, the artificial *β*-sheet forming proteins significantly inhibited the induction of the HSR in MG132-treated HEK293 cells (monitored using a Fluc reporter downstream of the *Hspa1a* promoter) (Olzscha, Schermann et al. 2011). Taken together, these findings suggest a possible common underlying mechanism in neurodegenerative diseases, whereby disease-associated proteins evade or attenuate the HSR, which facilitates inclusion formation and propagation throughout the CNS. Future research that incorporates a range of wild-type and disease-causing, aggregation-prone proteins could establish whether this is a molecular pathology common to all neurodegenerative diseases.

We hypothesised that the rate of protein aggregation or the surface area of the inclusion exposed to the cytosol would play a significant role in inducing the HSR. Therefore, flow cytometry and live cell confocal imaging experiments were performed to assess whether the capacity of cells to induce an HSR could be attributed to: (i) the rate of inclusion formation of the protein, (ii) the physical properties of the inclusions formed (i.e. size and granularity), and/or (iii) the intracellular concentration of the aggregation-prone protein. To our knowledge, this is the first study to report on the effects of aggregate size and rate of aggregate formation on the induction of the HSR. Since cells expressing Htt^72Q^ exhibited lower HSR induction compared to Fluc^DM^, we hypothesised that the rate of inclusion formation could be a key determinant in induction of the HSR. Interestingly, there was no significant difference in the rate at which Htt^72Q^ and Fluc^DM^ formed inclusions. Thus, the rate of inclusion formation does not influence HSR induction in this cell-based model. Another factor possibly influencing HSR induction is the size and granularity of the inclusions formed by each protein, since this would be an indirect measure of the surface area available to interact with cytoplasmic proteins, including regulators of the HSR. The recent development of FloIT provided an avenue to assess this as it enables the size and granularity of inclusions to be determined (Whiten, San Gil et al. 2016). Interestingly, the SOD1^WT^ and Htt^25Q^ aggregates detected demonstrated significantly greater granularity compared to the other proteins tested. Fluc^DM^ formed inclusions that were significantly smaller than those formed by Htt^72Q^ but were similar in size those formed by SOD1^G93A^. Since SOD1^G93A^ only induced an HSR in a low proportion of cells, these findings indicate that the size and granularity of the aggregate does not influence induction of the HSR in these cells.

High intracellular levels of proteins that exceed their predicted solubility is a key determinant of protein aggregation. This “supersaturation” concept is thought to be a significant factor driving protein aggregation in neurodegenerative diseases (Ciryam, Kundra et al. 2015). Indeed, proteins associated with ALS, such as TDP-43, FUS and SOD1 are supersaturated in the cell and, in their wild-type form, are susceptible to destabilisation and aggregation under conditions that drive proteostasis imbalance (Ciryam, Lambert-Smith et al. 2017). This instability is exacerbated by familial mutations in proteins linked to neurodegenerative diseases resulting in inherently aggregation-prone, supersaturated proteins in inherited neurodegenerative diseases (Yerbury, Ooi et al. 2019). Using our Neuro-2a (HSE:EGFP) cells we determined whether increasing concentrations of aggregation-prone proteins (and thus their level of saturation) resulted in induction of the HSR. Our data show that there is a strong positive correlation between the amount of the aggregation-prone protein in cells (SOD1^G93A^, Htt^72Q^, Fluc^DM^ and Fluc^WT^) and HSR induction. Significantly, this correlation was not observed for non-pathogenic forms of these disease-related proteins which are much less prone to aggregation (SOD1^WT^ and Htt^25Q^). Thus, the levels of these proteins *per se* does not drive HSR induction; rather, it is the susceptibility of the protein to aggregation combined with high intracellular levels. Lower concentrations of Fluc^DM^ and Fluc^WT^ induced an HSR compared to Htt^72Q^ and SOD1^G93A^ and this could suggest that the form of aggregation (*i.e.* amorphous versus amyloidogenic) may also be an important factor in HSR induction. HSR was more sensitive to increasing levels of Fluc^DM^ and Fluc^WT^, and/or the induction of the HSR was impaired/evaded by Htt^72Q^ and SOD1^G93A^ until a critical concentration was reached. These findings suggest that with increasing saturation of the protein there was an attempt by the cell to restore proteostasis after the accumulation of aggregation-prone proteins by inducing an HSR. Future research could work towards understanding whether the HSR is triggered in response to high concentrations of the soluble aggregation-prone protein or aggregated forms of protein.

In summary, this work shows that neurodegenerative disease-associated proteins are poor inducers of the HSR. Extracellular protein aggregates fail to induce the HSR in neuronal-like cells. Most remarkably, the intracellular expression of pathogenic aggregation-prone proteins also has a limited capacity to induce an HSR. Uniquely, we were able to determine the effect of a number of different factors pertaining to protein aggregates on HSR induction, including the aggregation propensity of pathogenic and non-pathogenic proteins, protein concentration within cells, number of inclusions formed, physical properties of the inclusions (size and granularity), and rate at which inclusions formed. Based on flow cytometric and live-cell imaging data it is concluded that HSR induction is dependent on the susceptibility of the protein to aggregation and the levels of the aggregation-prone protein in cells. Induction of the HSR was not affected by the size of the inclusions nor the rate of inclusion formation. The limited capacity of disease-related protein aggregation to induce an HSR suggests that these evade detection by the pathway leading to activation of the HSR and/or impair the HSR, however, the mechanism by which this occurs is yet to be elucidated. Our work also suggests there is therapeutic potential in the development of approaches that activate the HSR, and hence increase the levels of stress-response proteins including molecular chaperones, in order to inhibit further protein aggregation and promote cell viability in the context of neurodegenerative diseases.

## Author contributions

RSG and HE formulated the experimental approach. RSG performed all the experiments, analysed the data, constructed the figures and wrote the initial manuscript. DC and LM made the recombinant α-synuclein and SOD1^G93A^ protein and provided the protocols used for the *in vitro* aggregation of these proteins. RSG, DC, LM, AKW, JJY, LO, and HE edited the manuscript and approved the submission of the final manuscript.

## Acknowledgements

This research performed by RSG has been conducted with the support of the Australian Government Research Training Program Scholarship. We would like to thank Dr David Mitchell from the Australian Institute of Innovative Materials (University of Wollongong Australia) for his help with transmission electron microscopy. We thank the Illawarra Health and Medical Research Institute for technical and administrative support.

