## Supplementary Data for "Neurodegenerative disease-associated protein aggregates are poor inducers of the heat shock response in neuronal-like cells"

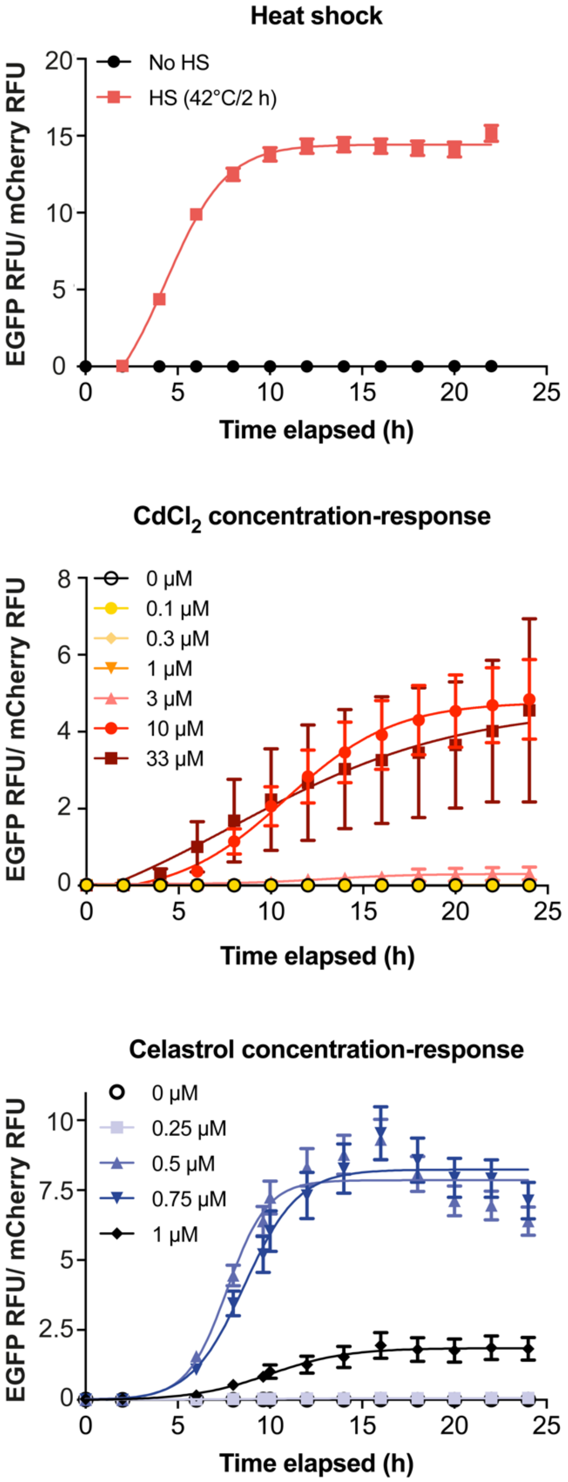


**Supplementary Figure 1. Monitoring induction of the heat shock response in Neuro-2a (HSE:EGFP) cells following heat shock, or treatment with $CdCl_2$ or celastrol .** Cells were subjected to heat shock (42°C/ 2 h), or log and ½ log doses of $CdCl_2$ (0-33 µM) or celastrol (0-1 µM). Following treatment cells were imaged every 2 h using an IncuCyte Live Cell Imaging System to monitor the levels of EGFP as a measure of HSR induction. Data is presented as the fold change in EGFP RFU over time, normalised to mCherry RFU, to account for changes in cell density over the time-course of the experiment. The data shown is the mean ± SEM of three independent repeats.

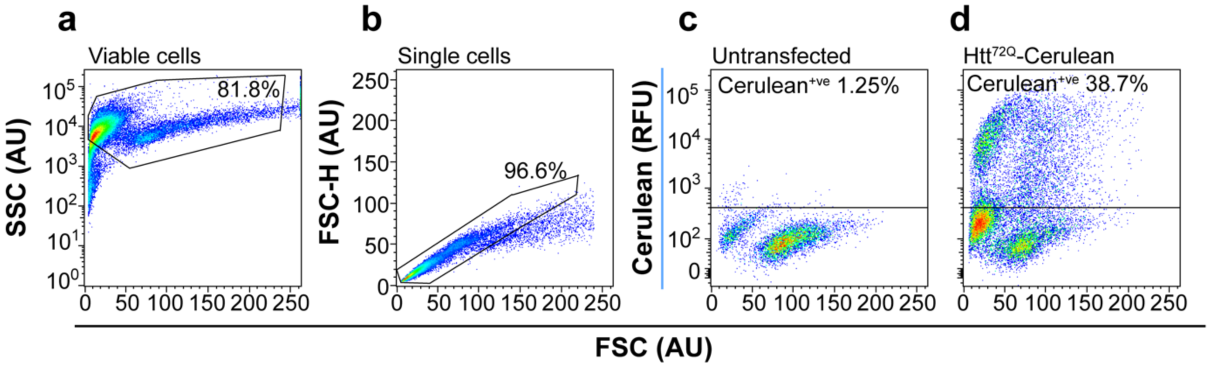


**Supplementary Figure 2. Gating strategy adopted for flow cytometric analysis of HSR induction in Neuro-2a (HSE:EGFP).** Neuro-2a cells were transfected to express cerulean tagged $SOD1^{WT}$, $SOD1^{G93A}$, $Htt^{25Q}$, $Htt^{72Q}$, $Fluc^{WT}$, or $Fluc^{DM}$ proteins and, 48 h post-transfection, cells were analysed by flow cytometry. (a) Plots of FSC and SSC of cells were used to resolve cellular debris and cell clumps. A polygonal gate was set around cells of interest and all downstream analyses was performed on this population. (b) Plots of FSC-height (FSC-H) and FSC-area (FSC-A) were used to resolve singlet and doublet events and a polygonal gate was used to exclude all doublets from downstream analyses. (c) Untransfected cells were used as a $cerulean^{-ve}$ control to identify $cerulean^{+ve}$ events. (d) Representative plot of Neuro-2a (HSE:EGFP) cells transfected to express $Htt^{72Q}$-cerulean are shown and the percent of $cerulean^{+ve}$ cells in the gate are denoted. Data shown are representative of three independent repeats.

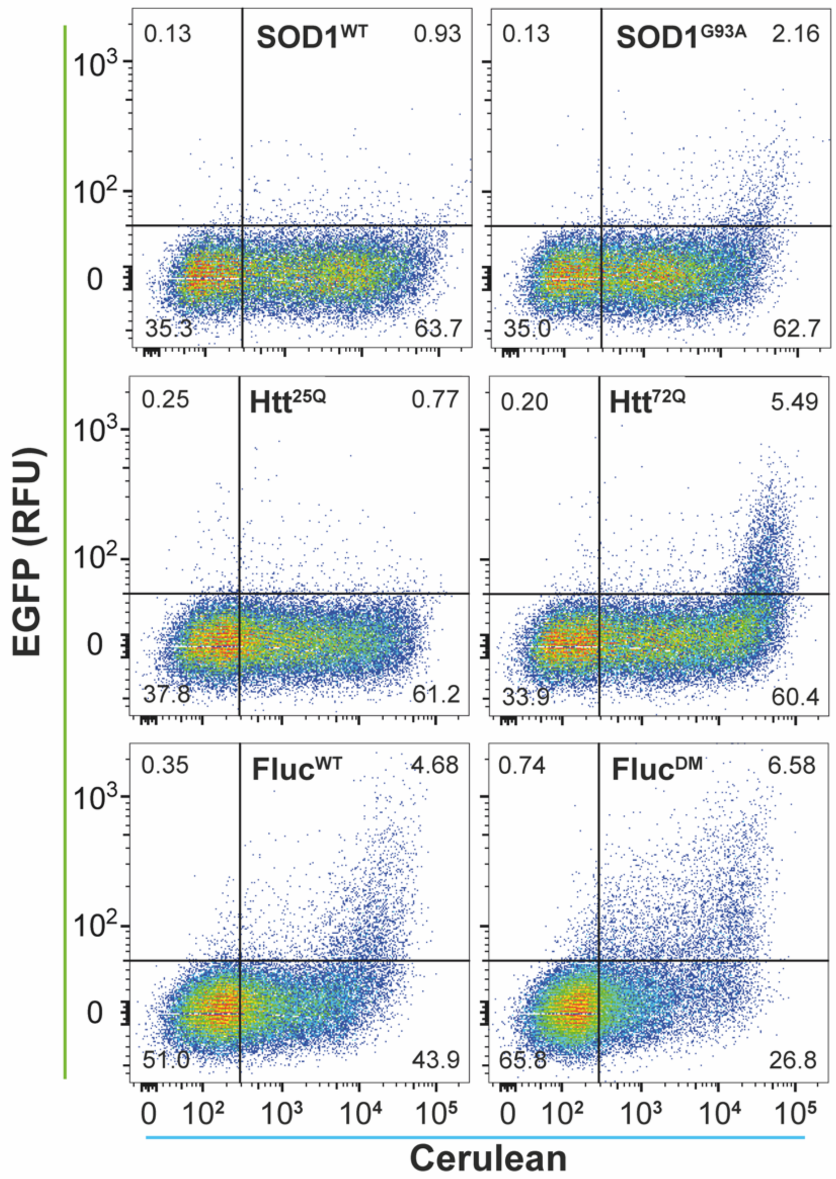


**Supplementary Figure 3. The HSR is differentially induced in response to the over-expression of pathogenic and non-pathogenic proteins.** Neuro-2a (HSE:EGFP) cells were transfected to express cerulean tagged $SOD1^{WT}$, $SOD1^{G93A}$, $Htt^{25Q}$, $Htt^{72Q}$, $Fluc^{WT}$ or $Fluc^{DM}$ proteins. After 48 h incubation, cells were harvested for analysis by flow cytometry. Cellular debris, cell clumps, doublet events and untransfected cells were excluded from the analysis as described in Supplementary Figure 1. Representative flow cytograms are shown for each of the cerulean-tagged proteins over-expressed in Neuro-2a (HSE:EGFP) cells. The proportion of cells in each quadrant gate is shown.

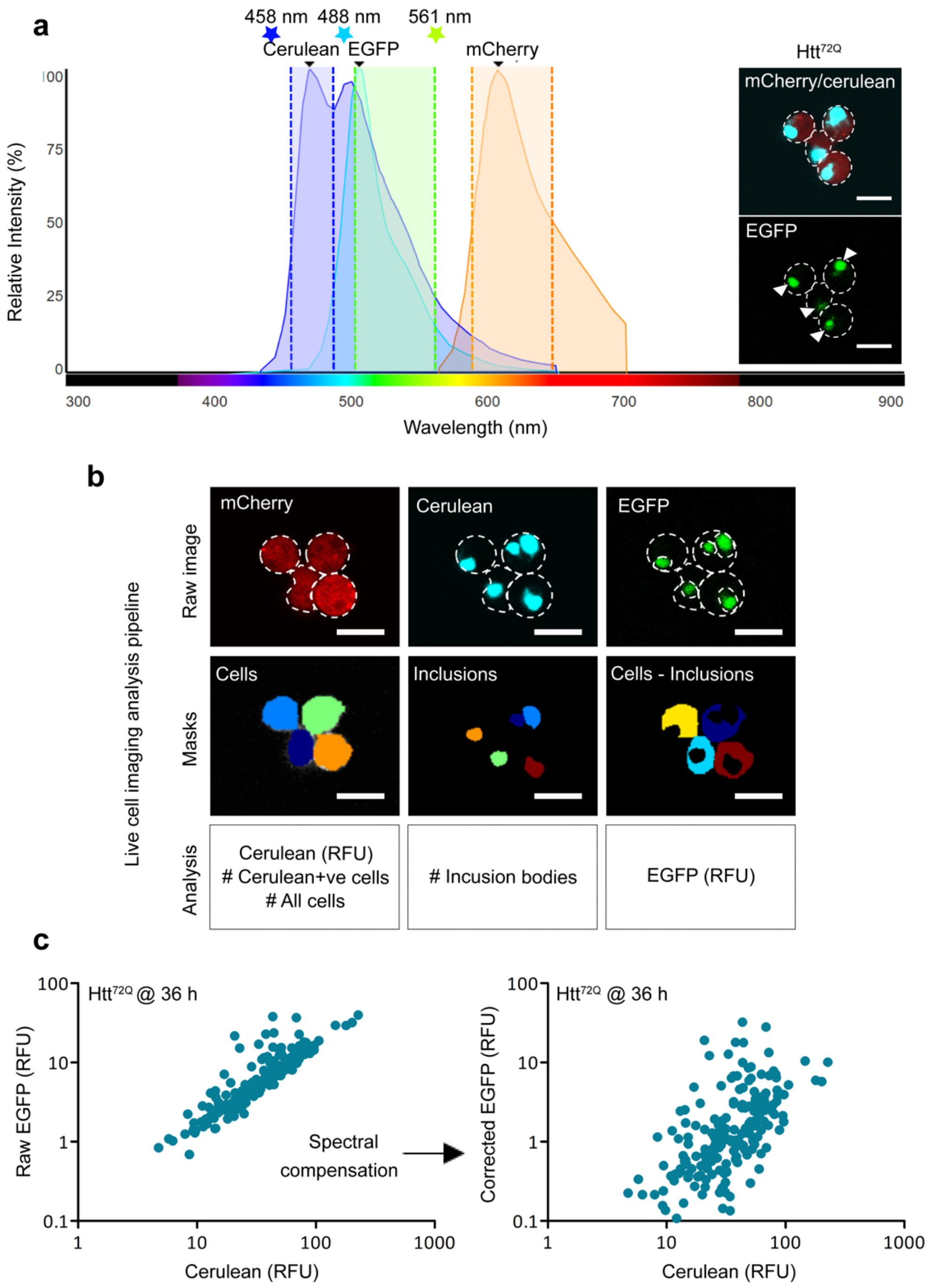


**d**

$$Raw\ EGFP - \left(\frac{Raw\ Cerulean}{1} \times \frac{13}{100}\right) = Corrected\ EGFP$$

**Supplementary Figure 4. Experimental design and analysis of live cell imaging experiments.** Neuro-2a (HSE:EGFP) cells were transfected to express Htt$^{72Q}$ or Fluc$^{DM}$ and imaged over time by confocal microscopy. (a) Fluorescence emission spectra of cerulean, EGFP and mCherry fluorescent proteins. The windows for emission collection were set at 462-492 nm (blue dotted lines), 506-563 nm (green dotted lines), and 600-657 nm (orange dotted lines), for cerulean, EGFP and mCherry, respectively. The lasers used to excite each fluorescent protein are depicted by the coloured stars. Inset: A representative image of Neuro-2a (HSE:EGFP) cells transfected to express Htt$^{72Q}$, demonstrating spectral overlap between the cerulean and EGFP channels (white arrowheads). (b) Analysis pipeline used in Cell

Profiler of confocal images acquired during the live cell imaging experiment. The mCherry channel (*left*) was used to define the "cells", which are masked in different colours, and this region was used to measure cerulean fluorescence intensity, the number of cerulean$^{+ve}$ cells, and the total number of all cells in each image over the time-course. The cerulean channel (*middle*) was used to identify "inclusions" and this region was used to count the number of inclusions formed in each image over the time-course. The "inclusions" were subtracted from the "cells" to generate a third region for measurement defined as "cells – inclusions", and this region was used to determine the EGFP fluorescence intensity in this channel. All scale bars = 20 μm. (c) Bivariate plots of cerulean and EGFP fluorescence intensities from the "cell – inclusions" region in Htt$^{72Q}$ transfected cells at 36 h, before (left) and after (right) spectral compensation. (d) Equation used to apply spectral compensation on the EGFP data.
